# Laying Hens Affected By Newcastle Virus And Laryngotracheitis Virus Show Rapid Recovery After Treatment With Ivermectin

**DOI:** 10.1101/2020.07.29.226027

**Authors:** Manolo Fernandez-Díaz, Luis Guevara, Eliana Icochea, Angela Montalván, Doris Villanueva-Pérez, Manolo Fernandez-Sánchez, Julio Ticona, Mirko Zimic, COVID-19 Working Group in Perú

**Affiliations:** Laboratorios de Investigación y Desarrollo, FARVET; Laboratorio de Patología Aviar, Facultad de Medicina Veterinaria. Universidad Nacional Mayor de San Marcos; Laboratorio de Bioinformática, Biología Molecular y Desarrollos Tecnológicos. Facultad de Ciencias y Filosofía. Universidad Peruana Cayetano Heredia

## Abstract

This is an anecdotal observation of an intervention study involving laying hens from a commercial farm in the city of Chincha, Peru, who suffered an outbreak caused by Newcastle disease virus (NDV) and infectious laryngotracheitis virus (ILTV), confirmed by clinical observations, serological and molecular tests (PCR).

In addition to receiving standard treatment appropriate to the state of health at the time of the operation, a group of birds were treated with a single dose of ivermectin administered subcutaneously (0.2 mL of a 1% solution equivalent to 200 µg/kg body weight), with a group of control birds not receiving the treatment being reserved.

The results showed a remarkable recovery of symptoms after 24 hours of treatment among the birds that received ivermectin. At 4 days after treatment, the birds that received ivermectin showed visibly greater mobility and vivacity, as well as a recovery in egg production. PCR tests after 4 days of treatment with ivermectin were negative for NDV and ILTV.

These results are interesting and suggest a possible effect of ivermectin against NDV and ILTV in birds. More controlled studies are needed to confirm this hypothesis.

## INTRODUCTION

Newcastle disease is caused by an Orthoavulavirus type 1 (ICTV 2019) whose viscerotropic velogenic strains cause great economic losses to the poultry industry worldwide. In Peru, Newcastle velogenic viruses, genotype XII, have been identified as causing outbreaks in fighting birds and small flocks of broilers and commercial layers (Chumbe et al., 2017) (Lupe et al., 2015).

Infectious laryngotracheitis is a highly infectious avian respiratory disease, caused by Gallid herpes virus type I, known as infectious laryngotracheitis virus (ILTV) that affects, economically, the global poultry industry (Garcia M. and Spatz S. 2016). In Peru, ILTV has been associated with a 16% decrease in egg production, and a 18% increase in layer chicken mortality (Alvarado et al., 2013), and a 27.3% decrease in productive efficiency in broilers (Lupe et al., 2015).

Both diseases are also endemic in fighting birds, causing a considerable negative economic impact on this activity and representing a risk to the Peruvian industry.

In recent years, despite intensive poultry vaccination programs, these infections continue to be a serious problem and although mortality is not high, low productivity and physical alterations in egg quality cause very large economic losses.

For both infections, prevention is based on the monitoring of strict biosecurity measures, as well as the use of attenuated, inactivated or vectorized vaccines (Miller P.J. and Koch G., 2016) (Garcia M and Spatz S, 2016). Despite the sustained use of vaccination programs, it is common to identify repeated outbreaks, not only in fighting birds, but also in medium and small commercial broiler and layer farms, putting the poultry industry and the country’s economy at risk.

Ivermectin (IVM), discovered in 1976, is a drug that belongs to the lactated macrocyclic family, and since 1981 it has been used in veterinary medicine. It has a broad spectrum of activity against a number of nematodes, microfilariae, external parasites in domestic species (Jozef Vercruysse, 2002) and exotic birds (Thomas-Baker, 1986) and has a wide safety margin (Nguyen, 2018).

Previous studies have shown that ivermectin possesses *in vitro* antiviral activity against RNA and DNA viruses (Crump, 2017). Recent studies have shown evidence of a positive effect of ivermectin against SARS-CoV-2 causing COVID19, at the in-vitro level associated with high concentrations of the drug (Caly L et al., 2020). In another study, through *in vitro* cytotoxicity assays using MTT (3-(4,5-Dimethylthiazol-2-yl)-2,5-Diphenyltetrazolium Bromide) on primary lines of Newcastle virus (NDV)-infected chicken fibroblasts, and in a concentration range between 6.25-200 µg/mL, demonstrated potent antiviral activity of Ivermectin against NDV (Azeem et al., 2015).

Nguyen in 2018 used it indirectly as an endectocide to control West Nile virus (Nguyen, 2018).

To date, ivermectin has not been shown to possess direct antiviral activity in birds under rearing conditions, and is effective in reducing clinical symptoms and lesions in birds affected with infections caused by Newcastle disease virus and infectious laryngotracheitis virus (ILV).

In the present study, we reported the use of ivermectin in egg-producing hens, infected with NDV and ILTV, from a poultry farm located in Chincha Province, Ica, Peru. The treatment with ivermectin resulted in a fast recovery of symptoms of the animals.

## METHODS

### Study population

This report includes ten thousand laying hens from a poultry farm located in Chincha, Ica, Peru, which showed clinical symptoms suggestive of infection with velogenic Newcastle disease virus (NDV) and avian infectious laryngotracheitis virus (ILTV). A flock of 5,000 hens was affected with velogenic NDV and a second flock of 5,000 hens (from another chicken house), was coinfected by both NDV and ILTV.

According to the clinical symptoms and the classic pathology, the birds showed clinical signs, characteristic of laryngotracheitis and Newcastle disease (Figure 1A).

**Figure 1.**
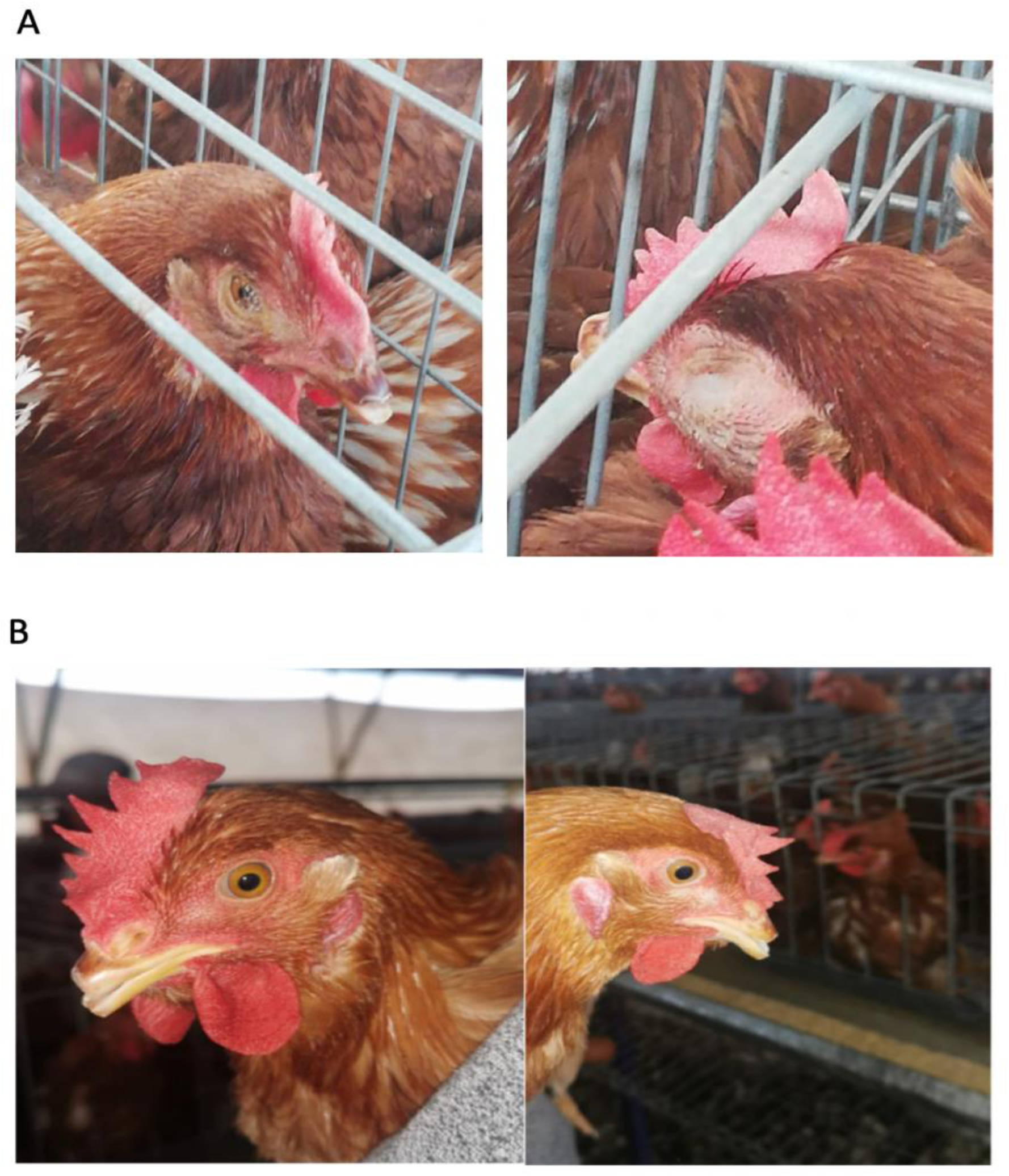
(A) Birds showing characteristic signs of laryngotracheitis such as depression and conjunctivitis at the day of treatment with ivermectin. (B) After 24 hours of treatment with ivermectin, clinical signs notably disappeared the birds

In addition to the clinical examination, the diagnosis of viral infection was confirmed by immunological tests for detection of specific antibodies, using the enzyme immunoassay (ELISA), as well as the molecular real-time PCR test to confirm the viral presence. A notable and significant drop in egg production was observed in these animals. A sample of 5 birds visibly affected was randomly selected from each of the two chicken houses at the day of treatment, for serology and molecular testing

### Diagnosis

#### Serology

The detection of antibodies against the ILTV was performed by ELISA test, using a commercial antibody detection kit from Biochek laboratories, USA, following the manufacturer’s recommendations. The inactivated viral antigen was attached to the plates for titration. The conjugate used was an alkaline phosphatase labelled anti-chicken antibody, and the chromogenic substrate was p-nitrophenylphosphate. The enzymatic reaction was stopped with buffered sodium hydroxide and the reaction was read at an absorbance of 405 nm.

To detect antibodies against the NDV virus, a commercial kit for the detection of antibodies from IDEXX laboratories was used following the manufacturer’s recommendations The viral antigen was coated in the 96-well plastic titration wells, then the serum to be investigated was placed and an anti-chicken serum, made from peroxidase conjugated goat and read at an absorbance of 650 nm, was used. The positivity of the sample was determined by the relationship between the value of the sample and the mean of the positive control.

#### Molecular detection

The real-time PCR technique was performed for NDV detection using Luna Universal One-Step RT-qPCR Kit E3005E (New England BioLabs, USA) following the manufacture procedures. Similarly, to detect ILTV virus, it was performed following the instructions of Luna Universal qPCR Master Mix Kit M3003E (New England BioLabs, USA). Samples of eyelids, trachea and cecal tonsils, were analyzed for the presence of NDV, ILTV, IBV, and aMPV viruses.

### Treatment

For the treatment of birds, ivermectin for parenteral use from Montana Peruvian Laboratories was used in all cases. 4,750 hens in the hen house affected only with NDV, received a multivitamin supplement comprising vitamins C, D and E. Within this same hen house, a row of 125 hens received only one dose of ivermectin subcutaneously (0.2 mL at 1%, corresponding to a final concentration of approximately 200 µg/kg body weight), and a row of 125 hens received no treatment at all (control group). Considering the critical health status, all the 5,000 hens in the second hen house affected with co-infection by NDV and ILTV, received the multivitamin supplement that includes vitamins C, D and E, as well as ivermectin subcutaneously (0.2 mL at 1%). At 4 days after treatment, 4 hens were randomly selected from the row that received ivermectin, and 4 hens from the control group row from the hen house affected only with NDV.

## RESULTS

### Diagnosis

At the time of intervention with ivermectin, high levels of antibody titers were obtained by the ELISA test in both flocks. This strongly suggested exposure to ILTV and NDV viruses. The molecular tests confirmed the presence of the ILTV, which was detected in eyelids and trachea. A velogenic NDV strain was detected in tracheas and cecal tonsils. No evidence of VBI or aMPV was found in any of the organs analyzed.

### Effect of treatment

At the time of the intervention, no mortality had been detected in the 2 chicken houses. After 24 hours, in most of the ivermectin treated hens, the symptoms disappeared for both the NDV and the NDV+ILTV affected animals. It was notable to observe that there were no clinical signs of avian infectious laryngotracheitis (Figure 1B). In contrast, the untreated group exhibited conjunctivitis (Figure 1A) and dyspnea. At 4 days after treatment, the ivermectin treated birds, showed clinically healthy birds with clean eyes compared to untreated birds, and no evidence of viral DNA was found in the molecular tests. In addition, the birds treated with ivermectin showed visibly greater mobility and vivacity (<u>VIDEO</u>)

Regarding egg production, after 3 days of treatment, birds that received ivermectin showed a slightly higher percentage of egg laying (66.01%) compared to birds that did not receive ivermectin (61.90%) (P= 0.15, Proportion test). After 6 days of treatment, birds that received ivermectin showed a higher percentage of egg laying (67.73%) compared to birds that did not receive ivermectin (61.88%) (P= 0.07, Proportion test). After 7 days, egg production returned to a normal egg-laying pattern in the animals treated with ivermectin, and showed a significantly higher percentage of egg laying (71.98%) compared to birds that did not receive ivermectin (64.68%) (P= 0.03, Proportion test).

After 4 days of treatment, in the flock of the birds affected by NDV and ILTV, a mortality of approximately 7 hens per day (out of 5,000 hens) was detected for 5 days. Necropsy showed that these 35 hens died with peritonitis typical of Newcastle disease due to egg rupture. In all cases, the organs of the respiratory tract (trachea and lungs) showed no macroscopic lesions. After 2 weeks, the birds affected by NDV and ILTV returned to their normal behavior.

## DISCUSSION

The present anecdotal report shows an interesting evidence of the efficiency of a single dose of Ivermectin, in the treatment and short recovery time of laying hens infected with avian NDV and ILTV. Remarkably, recovery of clinical signs for both diseases was achieved in less than 24 hours. Similarly, a recovery of egg production was shown in treated birds in a shorter time than usual (2 weeks versus 8 weeks that usually takes recovery). The dose used to control the viral infection in this case report, was 0.2 mL of a 1% ivermectin solution equivalent to 200 µg/kg body weight.

The mortality observed 4 days after treatment was very low (35/5,000), necropsies confirmed that chickens died from peritonitis associated with the rupture of eggs possibly caused by a previous fever.

The clinical, pathological and serological results were correlated with undetectable levels of viral load. These results confirm that the treatment was in some way effective in eliminating the virus, and not only in eliminating symptoms.

Several studies have shown evidence that ivermectin would have a potent antiviral action against HIV-1 and dengue virus, both dependent on the superfamily of importins which are necessary for different important metabolic processes of the cell (Caly et al., 2020). Ivermectin is suspected to play an important role in altering HIV-1 integrase and NS-5 polymerase (non-structural protein 5) in dengue virus (Wagstaff et al., 2015) (Yang et al., 2020). Evidence has been shown that ivermectin inhibits *in vitro* replication of some single-stranded RNA viruses such as Zika (Barrows et al., 2016), yellow fever and other alpha viruses (Varghese et al., 2016) (Mastrangelo et al., 2012). Thomas reported the short-term elimination of nematodes that parasitized the ocular nictitating membrane of exotic birds in a US zoo (Thomas-Baker, 1986). Recent studies suggest that Ivermectin is a potent specific inhibitor of import-mediated nuclear transport α/β, showing antiviral activity against several RNA viruses, blocking specific pathways of nuclear traffic in viral proteins (Caly L et al., 2020)

Some characteristics of ivermectin comprise, a low solubility, strong binding and rapid decomposition in the soil, rapid degradation under sunlight, low bioconcentration factor, low plant absorption, and it does not spread with percolation water, nor does it accumulate in the environment. In addition, at high levels, it has not been detected in animal wastes at toxic levels, nor in most plants and animals, terrestrial or aquatic, so it would not cause adverse environmental effects (Bruce et al., 2000).

Avian infectious laryngotracheitis in the commercial layers causes serious damage to the economy of the world’s small and large farms (Lupe López et al., 2015) (Colvero et al., 2015) (Sadiq and Mohammed, 2017). Despite vaccines, due to the ubiquity of the virus due to latency at the trigeminal nerve level, the infection is endemic (García M and Spatz S, 2016). Similarly, the velogenic and viserotropic strains of the Newcastle disease virus circulate in many parts of the world, having a major impact not only on the poultry industry, but also on farmed poultry, backyard birds, and fighting birds. The production of poultry meat and poultry eggs constitutes the source of income and food for many poor rural populations in the world (Terfa ZG et al., 2018), however, Newcastle disease is the main health threat that affects this type of bird. Therefore, it is necessary to have cost-effective drugs that inactivate or inhibit the replication of this virus.

In conclusion, the evidence presented here suggests that avian viral infections caused by NDV and ILTV could be controlled with ivermectin. Appropriate controlled studies that include a challenge to the birds under treatment are needed to support this hypothesis.

